# asymptoticMK: A web-based tool for the asymptotic McDonald–Kreitman test

**DOI:** 10.1101/101808

**Authors:** Benjamin C. Haller, Philipp W. Messer

## Abstract

The McDonald–Kreitman (MK) test is a widely used method for quantifying the role of positive selection in molecular evolution. One key shortcoming of this test lies in its sensitivity to the presence of slightly deleterious mutations, which can severely bias its estimates. An asymptotic version of the MK test was recently introduced that addresses this problem by evaluating polymorphism levels for different mutation frequencies separately, and then extrapolating a function fitted to that data. Here we present asymptoticMK, a web-based implementation of this asymptotic McDonald–Kreitman test. Our web service provides a simple R-based interface into which the user can upload the required data (polymorphism and divergence data for the genomic test region and a neutrally evolving reference region). The web service then analyzes the data and provides plots of the test results. This service is free to use, open-source, and available at http://benhaller.com/messerlab/asymptoticMK.html.

## INTRODUCTION

The extent to which molecular evolution is driven by positive selection, rather than neutral evolutionary processes such as random genetic drift, is one of the central questions of modern evolutionary biology. This question can be studied quantitatively by estimating the parameter *α*, which specifies the fraction of nucleotide substitutions in a given genomic region that were driven to fixation by positive selection (Eyre-Walker 2006). Values of *α* close to one indicate that most substitutions in the region were indeed the result of positive selection, whereas values close to zero indicate neutral evolution.

One of the most widely used approaches for inferring *α* from polymorphism and divergence data is the McDonald–Kreitman (MK) test (McDonald and Kreitman 1991; Eyre-Walker 2006), which compares levels of divergence between a genomic test region and a neutrally evolving reference region with the levels of polymorphism in the two regions. Early applications of the MK test typically focused on nonsynonymous sites in protein-coding regions as the test region, while synonymous sites were used as the neutral reference. However, the approach can also be applied to arbitrary genomic compartments or classes of mutations (Andolfatto 2005).

The original MK test makes several critical assumptions about the nature of the evolutionary process. First, it assumes that the positively selected mutations that ultimately contribute to divergence in the test region go to fixation quickly, such that they do not contribute noticeably to polymorphism levels. Second, it assumes that deleterious mutations in the test region are sufficiently deleterious to be lost quickly, such that they contribute to neither polymorphism nor divergence. Finally, neutral mutations in the test region are assumed to be subject to drift similar to the mutations in the neutral reference region and can therefore contribute to both polymorphism and divergence. Under these assumptions, it holds that

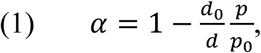

where *d* and *d*_0_ are the levels of divergence in the test region and neutral reference region, respectively, while *p* and *p*_0_ specify the respective levels of polymorphism in the two regions (Eyre-Walker 2006).

With the growing availability of genome-level polymorphism and divergence datasets, the MK test has become a popular method for inferring positive selection in various organisms (Fay 2011). Several software tools and web services with implementations of the test have also been developed (Egea *et al*. 2008; Librado and Rozas 2009; Eyre-Walker and Keightley 2009; Stoletzki and Eyre-Walker 2011; Vos *et al*. 2013). The estimates of *α* obtained in these studies range from as high as ~0.5 for nonsynonymous substitutions in *Drosophila* (Sella *et al*. 2009), to close to zero in organisms such as yeast (Elyashiv et al. 2010) or many plants (Gossmann *et al*. 2010). Indeed, estimates of *α* obtained from Equation (1) are often negative, indicating that at least some of the assumptions of the test were likely not met.

One major problem with the original MK test lies in its assumption that deleterious mutations do not contribute to polymorphism in the test region. This stands in contrast to the frequent observation of weakly deleterious mutations in many organisms, and the fact that such mutations can substantially affect the site frequency spectrum (SFS) of polymorphisms in functional genomic regions (Bustamante *et al*. 2005; Eyre-Walker *et al*. 2006). In the presence of weakly deleterious mutations, *p* will overestimate the rate at which polymorphisms go to fixation in the test region, which will bias estimates of *α* downwards (providing one possible explanation for the frequent observation of negative *α* values).

As one strategy to address this problem, it has been proposed to only consider polymorphisms for which the derived allele is above a certain threshold frequency when estimating *p* and *p*_0_ (Charlesworth and Eyre-Walker 2008). This is because the fraction of weakly deleterious mutations among all polymorphisms should be lower for higher derived-allele frequencies. Ideally, one would wish to set this cutoff high, to minimize the bias due to weakly deleterious mutations; however, the higher this cutoff, the fewer polymorphisms will actually remain in the dataset, thus increasing statistical noise. To circumvent this problematic tradeoff, more sophisticated extensions of the original MK test first attempt to infer the actual distribution of fitness effects among new mutations in the test region from the SFS, and then correct fixation probabilities accordingly (Boyko *et al*. 2008; Eyre-Walker and Keightley 2009). Yet these approaches can still suffer from unknown effects of demography or linked selection that are also expected to affect the shape of the SFS. The most sophisticated extensions of the test therefore additionally incorporate basic demographic models to improve their estimates (Keightley and Eyre-Walker 2007; Boyko *et al*. 2008; Eyre-Walker and Keightley 2009), which requires additional (and often uncertain) assumptions about the demographic history of the population of interest.

In contrast to such model-based approaches, a considerably simpler, heuristic approach was recently proposed by Messer and Petrov (2013). This approach generalizes the frequency-cutoff approach described above, without the need to discard polymorphism data. Instead of setting a specific frequency cutoff, it separately estimates *α* for each of a set of discrete mutational frequency classes:
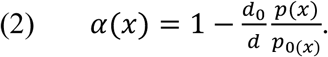

Here *p*(*x*) and *p*_0_(*x*) specify the levels of polymorphism in the test and reference regions, respectively, considering only those polymorphisms for which the derived allele is present at frequency *x* in the population (estimated from a population sample, for example). In the presence of deleterious mutations, *α*(*x*) will underestimate the true value of *α* for small *x*, yet should converge to the correct value as *x* approaches one. The asymptotic estimate of *α* is then obtained by fitting a function *α*_fit_(*x*) to the empirical *α*(*x*) values and extrapolating this function to *x* = 1:
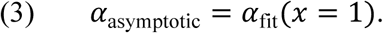

One key advantage of this approach is that because *α*(*x*) does not depend on the individual functions *p*(*x*) and *p*_0_(*x*) but only on their ratio, any biases due to demography or linked selection that affect the SFS in the test and reference regions in the same way will effectively cancel out (Messer and Petrov 2013). Another advantage over model-based approaches is that the asymptotic McDonald–Kreitman approach is much more computationally efficient, as it requires only fitting a simple curve to the data.

In this paper, we present asymptoticMK, a web-based tool for executing the asymptotic McDonald–Kreitman test quickly and easily in any web browser. After the necessary values are entered, asymptoticMK generates analyses and plots that are directly usable in publications. It is based internally on R, but no knowledge of R is needed to use it, nor does the user of asymptoticMK need to have R installed on their computer. For those who do wish to run the test themselves in R, the necessary code is freely available online. The asymptoticMK service can also be run in an automated fashion at the command line, for bulk analysis in script-based workflows.

## MATERIALS AND METHODS

### Implementation

The asymptoticMK web service is implemented in R (R Development Core Team 2016). It uses the package `FastRWeb` (Urbanek 2008) to parse HTTP requests and generate responses, and uses the package `Rserve` (Urbanek 2003) as the lower-level interface that communicates with the web server through the standard CGI mechanism.

### Usage

The web service is free to use, without license restrictions of any kind, and is available at http://benhaller.com/messerlab/asymptoticMK.html. That URL displays an entry page (Figure 1) with an input form in which the user may enter the necessary data for the test: *d* (the substitution rate in the test region), *d*_0_ (the substitution rate in the neutral reference region), and an uploaded file containing tab-delimited rows of data with values for *x* (the derived allele frequency), *p*(*x*) (the polymorphism level in the test region at that frequency), and *p*_0_(*x*) (the polymorphism level in the neutral reference region at that frequency). A sample polymorphism file is provided on the website. In practice, it is often advisable to combine polymorphism levels into a smaller number of frequency bins, where *x* then specifies the central frequency of the bin. This is particularly relevant when the data includes frequencies at which no polymorphisms are actually present in the neutral region, in which case *α*(*x*) would be undefined for those particular frequencies according to Equation (2). The input form also allows entry of minimum and maximum values defining a cutoff interval for *x*, such that the test is run using only the polymorphisms whose frequencies fall within that cutoff interval; this is usually desirable as a means of excluding the lowest‐ and highest-frequency polymorphisms, where SNP quality issues and polarization errors are generally most pronounced. This frequency cutoff is set to [0.1, 0.9] by default.

**Figure 1.**
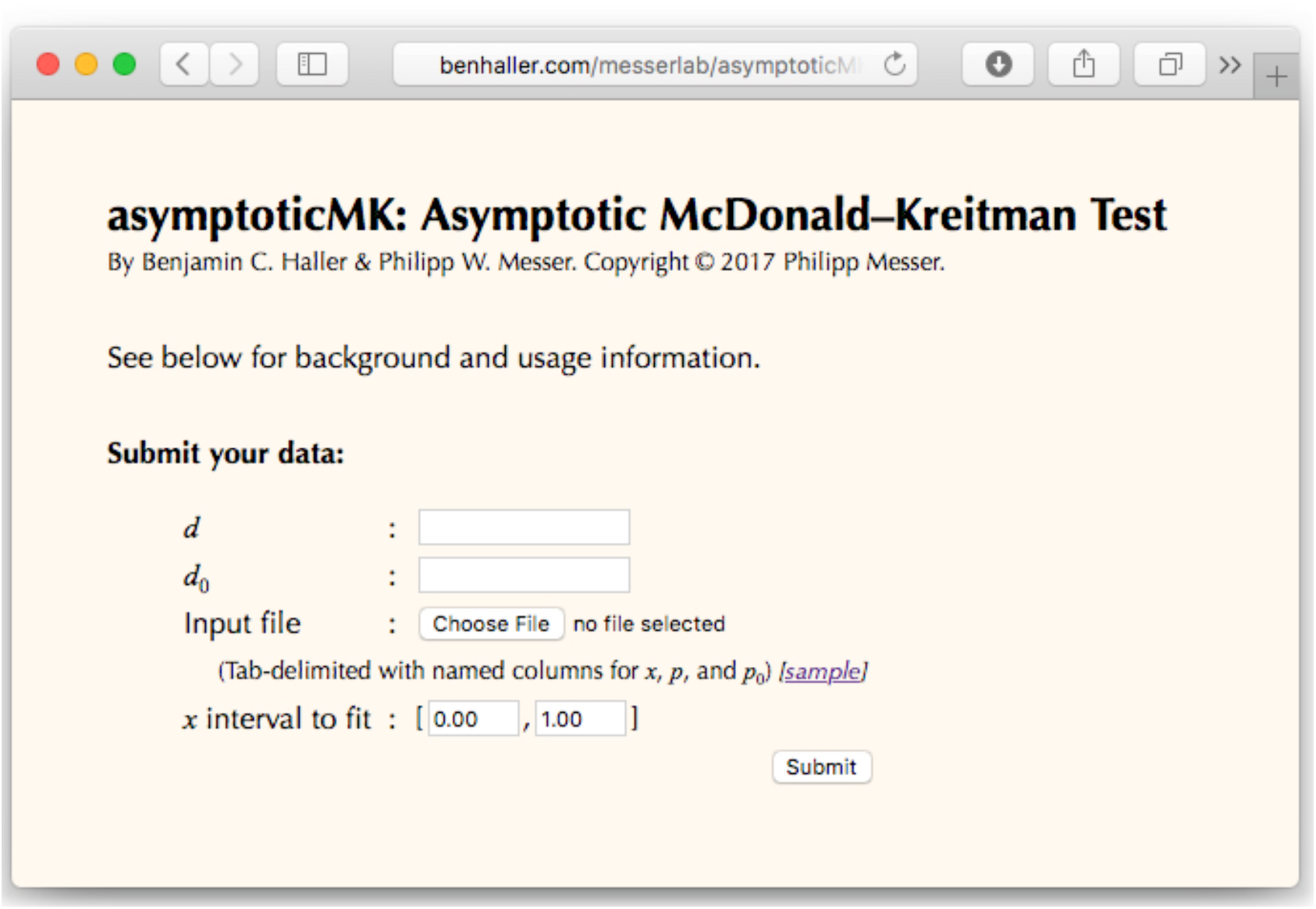
A screenshot of the web page for asymptoticMK. After entering values for *d* and *d*_0_, choosing an input file with binned values for *x*, *p*, and *p*_0_, and choosing the *x* interval to fit, the user can click the Submit button and asymptoticMK will provide its results in a new browser window or tab.

Upon submission of the web form, asymptoticMK conducts its analysis and then opens a results page in a new browser tab, presenting a summary of the input data and the results from the analysis. The first plot on this results page shows binned polymorphism counts, *p*_0_(*x*) and *p*(*x*), for the submitted data; the second plot shows that same data normalized (i.e., the normalized SFS in the test and reference regions). A third plot shows the calculated empirical *α*(*x*) as a function of *x*, estimated from the input data according to Equation (2). The fourth plot shows the same *α*(*x*) data, with the best-fitting model and the asymptotic estimate of *α* superimposed upon the data.

Below these plots, the results of the analysis are presented in two tables. The first table provides the coefficients *a*, *b*, and (for exponential fits) *c* of the model yielding the best fit to the data. The second table provides the estimated *α*_asymptotic_ according to Equation (3), and the upper and lower limits of the 95% confidence interval around that estimate, as well as the estimated *α* from the original non-asymptotic McDonald–Kreitman test (*α*_original_) for comparison (also estimated from all polymorphisms falling within the frequency cutoff interval specified on the input page).

For purposes of automation, asymptoticMK can also be run at the command line using the Linux/Unix `curl` command. For example, the command

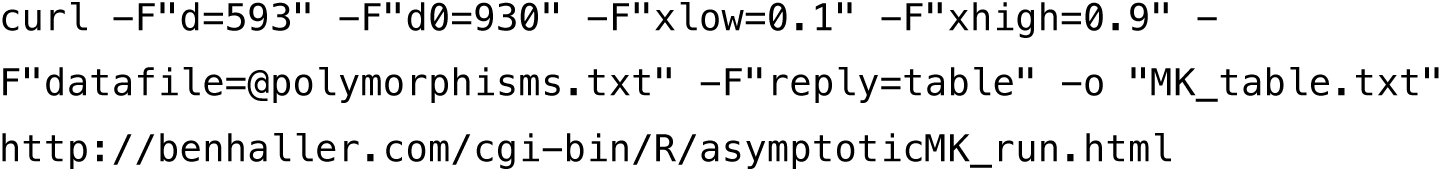

would run asymptoticMK with the given values of *d* and *d*_0_, the given *x* cutoff interval, and polymorphism data uploaded from the local file `polymorphisms.txt`, and would output a simple table of results to the file `MK_table.txt`. Further documentation on the use of this feature is provided on the asymptoticMK web page.

### Fitting and analysis procedure

The asymptotic McDonald–Kreitman test first involves calculating values of *α*(*x*) by applying Equation (2) to each frequency bin provided, as described by Messer and Petrov (2013). The next step involves fitting a function *α*_fit_(*x*) to these empirical *α*(*x*) values. For greater robustness, asymptoticMK fits two functions to the data. The first function is exponential, of the form *α*_fit_(*x*) = *a* + *b* exp(−*cx*) and is fitted using the `nls2()` function, from the R package `nls2` (Grothendieck 2013). This fit is done in two steps. First, a brute-force scan for the closest fit is conducted across the likely portion of the three-dimensional parameter space defined by *a*, *b*, and *c*, by exhaustive search. This supplies reasonably good starting values for the second step, which refines those starting values using standard nonlinear least-squares regression. This two-step procedure generally works well, but can occasionally fail to converge if the data is not, in fact, exponential in form.

To address this possibility of nonconvergence of the exponential fit, asymptoticMK also fits a linear function of the form *α*_fit_(*x*) = *a* + *bx*, with the `lm()` function that is part of the `stats` package included in R. This fit always converges, and thus provides a backstop that allows the test to complete even when given irregular or extremely noisy data; however, it is always recommended that the results of the analysis be inspected visually to confirm that they are in fact meaningful.

Once these two models have been fitted, asymptoticMK chooses which model will be used for the remainder of the analysis. If the exponential fit failed to converge, then the linear model is chosen; if both fits succeeded, then the better model is chosen using the Akaike information criterion (AIC). Occasionally, in pathological cases, the exponential fit will have the better AIC but will have extremely large coefficient standard error(s); in this case, the linear fit is chosen since predictions from the exponential model would be effectively worthless.

The chosen model is then used to provide an estimate of the value of *α*_asymptotic_ according to Equation (3), by evaluating the fitted function *α*_fit_(*x*) at *x* = 1; this is the primary result of the test, and provides the test’s estimate of the true value of *α* within the test region. A 95% confidence interval around this estimate is also calculated. For the exponential model, this is done using Monte Carlo simulation based upon the fitted model, using the `predictNLS()` function published online by Spiess (2013); for the linear model, it is done using the standard R function `predict()`.

### Test datasets

To provide a test of asymptoticMK using empirical data, we used the same *Drosophila melanogaster* dataset that Messer and Petrov (2013) used in their Figure 3C. This data set consists of SNPs obtained from the genome sequences of 162 inbred fly lines generated by the *Drosophila* genetic reference panel (Mackay *et al*. 2012). Divergence data was obtained from genome alignments between *D. melanogaster* and *D. simulans*, extracted from the 12 *Drosophila* genomes data (Clark *et al*. 2007). The test data in the asymptoticMK analysis (*d* and *p*) are genome-wide nonsynonymous mutations, while synonymous sites were used as the neutral reference (*d*_0_ and *p*_0_). The polymorphism data is available online at http://benhaller.com/messerlab/sample_polymorphism_levels.txt, with associated values *d* = 59570 and *d*_0_ = 159058. The default frequency cutoff interval of [0.1, 0.9] was used in the analysis of this dataset with asymptoticMK.

We also tested asymptoticMK on simulated data, using the forward genetic simulation framework SLiM 2 (Haller and Messer 2017). A population of 1000 diploid individuals was simulated to evolve for 200,000 generations. The simulated chromosome was 10^7^ base pairs long. Nucleotide mutations occurred uniformly at a rate of 10^−9^ per base per generation, and recombination occurred uniformly at a rate of 10^−7^ per base per generation. Each new mutation was either of neutral type “m1” (relative proportion of 0.5 of all new mutations), of functional non-beneficial type “m2” (relative proportion of 0.5 of all new mutations), or of functional beneficial type “m3” (a relative proportion of 0.0005 of all new mutations). The neutral m1 mutations always had a selection coefficient of *s* = 0.0; the selection coefficients of m2 mutations were drawn from a gamma distribution with a mean of *s* = −0.02 and a shape parameter of 0.2; and m3 mutations always had a selection coefficient of *s* = 0.1. Fitness effects were assumed to be codominant. A burn-in of 10000 generations was run to equilibrate the model. Every 500 generations thenceforth, all polymorphisms were recorded in the population by dividing them according to their frequency into 50 equal-width frequency bins, and then adding them to an ongoing binned tabulation. The SLiM configuration script used for these simulations is provided online at http://benhaller.com/messerlab/asymptoticMK_SLiM.html.

At the end of the model run, we obtained binned values for *p*(*x*) and *p*_0_(*x*), where *p*_0_ was estimated from all mutations of type m1, while *p* was estimated from the combined mutations of types m2 and m3. Values for *d* and *d*_0_ were obtained from the set of mutations fixed during the simulation; as with *p*_0_ and *p*, *d*_0_ was estimated from all mutations of type m1, while *d* was estimated from the combined mutations of types m2 and m3. These values, output by the model, were used in asymptoticMK with the default *x* interval of [0.1, 0.9]. The true value of *α* was also calculated by the SLiM model as the fraction *d*_3_ / (*d*_2_ + *d*_3_), where *d*_2_ is the number of m2 mutations fixed and *d*_3_ is the number of m3 mutations fixed. This value provides a metric for the accuracy of asymptoticMK – a benefit of using simulated data, where the true *α* can be calculated.

## RESULTS AND DISCUSSION

Results from our test of asymptoticMK with the empirical *D. melanogaster* dataset are shown in Figures 2A and 2B. The fitted exponential function is: *α*_fit_(*x*) = 0.585 − 0.622 exp(−3.80*x*). The asymptotic McDonald–Kreitman *α* estimate provided by this model is 0.571. These results match those obtained by Messer and Petrov (2013) using the same dataset (their Figure 3C), as expected. The *α* estimate provided by the original McDonald–Kreitman test is 0.407, by comparison (shown in Figure 2B).

**Figure 2.**
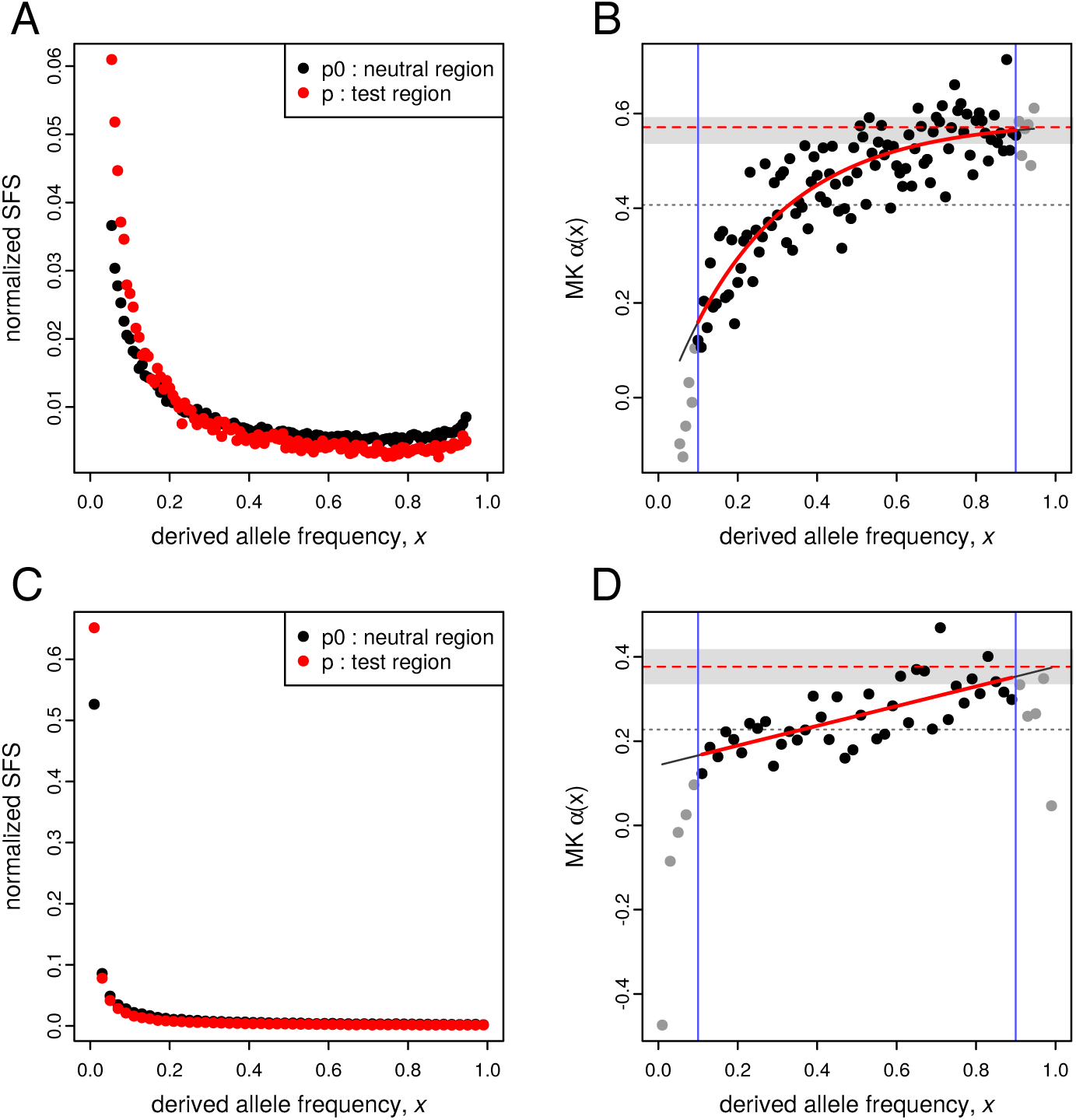
Results from asymptoticMK for two test datasets. A and B show results from the *Drosophila* dataset of Messer and Petrov (2013); C and D show results from a SLiM simulation run (see Materials & Methods). A and C show the normalized site frequency spectra (SFS) for their respective datasets. B and D show the results of the asymptotic MK test. In B and D, the two vertical blue lines show the limits of the frequency cutoff interval used for fitting. Points indicate binned values of *α*(*x*), estimated according to Equation 2; points are gray if they are outside the cutoff interval (and thus not used in fitting). The solid red curves show the fitted functions (exponential in B and linear in D). The dashed red lines show the estimates of *α*_asymptotic_, obtained from the fitted function according to Equation 3; this is the main result of the asymptotic MK test. The gray bands indicate the 95% confidence intervals around the *α*_asymptotic_ estimates. The dotted gray lines show *α*_original_, the estimates of *α* from the original (non-asymptotic) MK test, for comparison (also calculated using only the data within the cutoff interval).

The asymptoticMK results from the analysis of the SLiM simulation dataset are shown in Figures 2C and 2D. In this case, asymptoticMK deemed the linear fit to be superior to the exponential fit (and indeed, the data within the cutoff interval looks very linear in shape). The fitted linear function in this case is: *α*_fit_(*x*) = 0.143 + 0.234*x*. The asymptotic McDonald– Kreitman *α* estimate provided by this model is 0.377. This may be compared to the true *α* value provided by the simulation, 0.331. The *α* value from the original McDonald–Kreitman test within the cutoff interval, on the other hand, is 0.228 (shown in Figure 2D). If the cutoff interval is widened to [0.05, 0.95], the encompassed data then has a more exponential shape, and asymptoticMK then prefers the exponential fit (not shown), providing an improved asymptotic McDonald–Kreitman *α* estimate of 0.317 (as compared to the same true *α* value of 0.331, and an original McDonald–Kreitman *α* estimate of 0.170 within this cutoff interval). The default cutoff interval thus obscured the exponential shape of the data and prevented a good fit; this underlines the importance of choosing a cutoff interval that fits the shape of the data, and of examining the quality of the fit critically rather than simply accepting the default fit. Nevertheless, even the linear fit provided by the default cutoff interval provided a much closer estimation of the true *α* value than did the original non-asymptotic test.

In this paper, we presented asymptoticMK, a new web-based tool for executing the asymptotic McDonald–Kreitman test. To demonstrate its functionality, we analyzed both empirical data and a simulation-generated dataset. Results from both of these datasets illustrate the greater power of the asymptotic McDonald–Kreitman test to estimate the true value of *α*, compared to the estimates provided by the original non-asymptotic test. The asymptoticMK service presented here allows the user to obtain these results quickly and easily through any web browser.

## ACKNOWLEDGMENTS

The authors thank G. Grothendieck, J. Horner, and S. Urbanek for their contributions to the R packages used here. We also thank S. Urbanek for his unstinting help with the installation and use of `FastRWeb`, A.-N. Spiess for his `predictNLS()` function, and most of all the R Core Team for R itself. asymptoticMK is only possible because of the free software developed by these and other contributors. This work was supported by funds from the College of Agriculture and Life Sciences at Cornell University to PWM.

